# Integrative pathway enrichment analysis of multivariate omics data

**DOI:** 10.1101/399113

**Authors:** Marta Paczkowska, Jonathan Barenboim, Nardnisa Sintupisut, Natalie C. Fox, Helen Zhu, Diala Abd-Rabbo, PCAWG Network and Pathway Analysis Group, Paul C. Boutros, Jüri Reimand

**Author notes:** these authors contributed equally (Marta Paczkowska, Jonathan Barenboim).

## Abstract

Multi-omics datasets quantify complementary aspects of molecular biology and thus pose challenges to data interpretation and hypothesis generation. ActivePathways is an integrative method that discovers significantly enriched pathways across multiple omics datasets using a statistical data fusion approach, rationalizes contributing evidence and highlights associated genes. We demonstrate its utility by analyzing coding and non-coding mutations from 2,583 whole cancer genomes, revealing frequently mutated hallmark pathways and a long tail of known and putative cancer driver genes. We also studied prognostic molecular pathways in breast cancer subtypes by integrating genomic and transcriptomic features of tumors and tumor-adjacent cells and found significant associations with immune response processes and anti-apoptotic signaling pathways. ActivePathways is a versatile method that improves systems-level understanding of cellular organization in health and disease through integration of multiple molecular datasets and pathway annotations.

## Introduction

Pathway enrichment analysis is an essential step for interpreting high-throughput (*omics*) data that uses current knowledge of genes and biological processes. A common application determines statistical enrichment of molecular pathways, biological processes and other functional annotations in long lists of candidate genes^1,2^. Genomic, transcriptomic, proteomic and epigenomic experiments emphasize distinct and complementary aspects of underlying biology and are best analyzed integratively, as is now routinely done in large-scale projects such as The Cancer Genome Atlas (TCGA)^3^, Clinical Proteome Tumor Analysis Consortium (CPTAC), International Cancer Genome Consortium (ICGC)^4^, Genotype-Tissue Expression (GTEx)^5^ and others. Thus, simultaneous analysis of multiple candidate gene lists for characteristic pathways is increasingly needed. Numerous approaches are available for interpreting single gene lists. For example, the GSEA algorithm can detect up-and down-regulated pathways in gene expression datasets^6^. Web-based methods such as Panther^7^, ToppCluster^8^ and g:Profiler^9^ detect significantly enriched pathways amongst ranked or unranked gene lists and are generally applicable to genes and proteins from various analyses. Some approaches allow analysis of multiple input gene lists however these primarily rely on visualization rather than data integration to evaluate the contribution of distinct gene lists towards each detected pathway^8,9^. Finally, no methods are available for unified pathway analysis of coding and non-coding mutations from whole-genome sequencing (WGS) data, or integrating these with other types of DNA aberrations such as copy number changes and balanced genomic rearrangements. We report the development of the ActivePathways method that uses data fusion techniques to address the challenge of integrative pathway analysis of multi-omics data. We demonstrate the method by analyzing known and candidate cancer driver genes with coding and non-coding somatic mutations in 2,583 whole cancer genomes of the ICGC-TCGA PCAWG project^10,11^, prognostic pathways in breast cancer subtypes, and regulatory networks of tissue transcriptomes using the GTEx^5^ compendium.

Characterization of genes and somatic mutations that drive oncogenesis is a central goal of cancer genomics research. Cancer genomes are characterized by few frequently mutated pan-cancer drivers such as *TP53*, less-frequent drivers with primarily tissue-specific effects and numerous infrequently mutated genes often referred to as *the long tail*. The majority of currently known driver mutations affect protein-coding sequence^12^ and only few high-confidence non-coding drivers have been found, such as the mutation hotspots in the *TERT* promoter^13^. Discovery of non-coding driver mutations is a major goal of large cancer whole genome sequencing efforts such as PCAWG^10,11^. Pathway and network analysis of cancer mutations is a powerful approach that uses knowledge of coding driver genes and their pathway annotations as priors to assist in detection of weak driver variants including those in the non-coding genome^1^. The PCAWG project has produced a consensus dataset of predicted protein-coding driver genes (CDS) and non-coding regions of 5’ and 3’ untranslated elements (UTRs), promoters and enhancers of protein-coding genes across 2,583 whole cancer genomes of multiple cancer types^14^. Driver gene p-values in the dataset reflect the frequency and functional impact of somatic single nucleotide variants (SNVs) and small insertions-deletions (indels) in these protein-coding and non-coding genomic regions. Here we used our ActivePathways method to interpret these driver predictions with pathway information including biological processes of Gene Ontology^15^ and molecular pathways defined by Reactome^16^. Two further case studies focused on prognostic molecular pathways of breast cancer through integration of genomic and transcriptional alterations, and gene regulatory networks associated with organ growth control in healthy human tissues.

## Results

### Multi-omics pathway enrichment analysis with ActivePathways

ActivePathways is a simple four-step method that extends our earlier work^9^ (**Figure 1**). It requires two input datasets. First, a table of *gene p-values* contains multiple p-values for every gene representing different types of evidence such as gene significance in distinct omics experiments. These could include p-values evaluating the significance of differential gene expression in tissues of interest, gene essentiality, mutation or copy number alteration burden, and many others. Second, a collection of *gene sets* represents molecular pathways, biological processes and other gene annotations we refer to as *pathways*. Depending on the hypothesis, pathways may also include other types of gene sets such as targets of transcription factors or microRNAs. In the first step of ActivePathways, we derive an integrated gene list that aggregates significance from all types of evidence for each input gene. The integrated gene list is compiled by fusion of gene significance from different types of evidence using the Brown’s extension^17^ of the Fisher’s combined probability test, which conservatively adjusts for overall correlations of p-values in estimating the overall significance of every gene. The integrated input gene list is then ranked by decreasing significance and filtered using a lenient cut-off to capture a long tail of candidate genes and to filter the bulk of insignificant ones (unadjusted *P*_*gene*_<0.1). The integrated gene list is analyzed with a ranked hypergeometric test for each pathway to capture smaller pathways tightly associated with few top-ranking genes and broader processes with abundant albeit weaker signals from larger subsets of input genes. The stringent family-wise multiple testing correction method by Holm^18^ is applied across pathways to reduce false positives (*Q*_*pathway*_<0.05). In the third step, candidate gene lists corresponding to distinct types of evidence are separately evaluated using the above procedure. This step determines which pathways are significantly supported by each of the input omics datasets and also reveals corresponding genes in each pathway. Importantly, the step also highlights pathways that are only found through data integration and are not apparent in any single type of omics evidence alone. In the fourth step, the method provides input files for Enrichment Map^19^ for visualizing and reducing the redundant set of all detected pathways to a narrower, focused network of biological themes.

**Figure 1:**
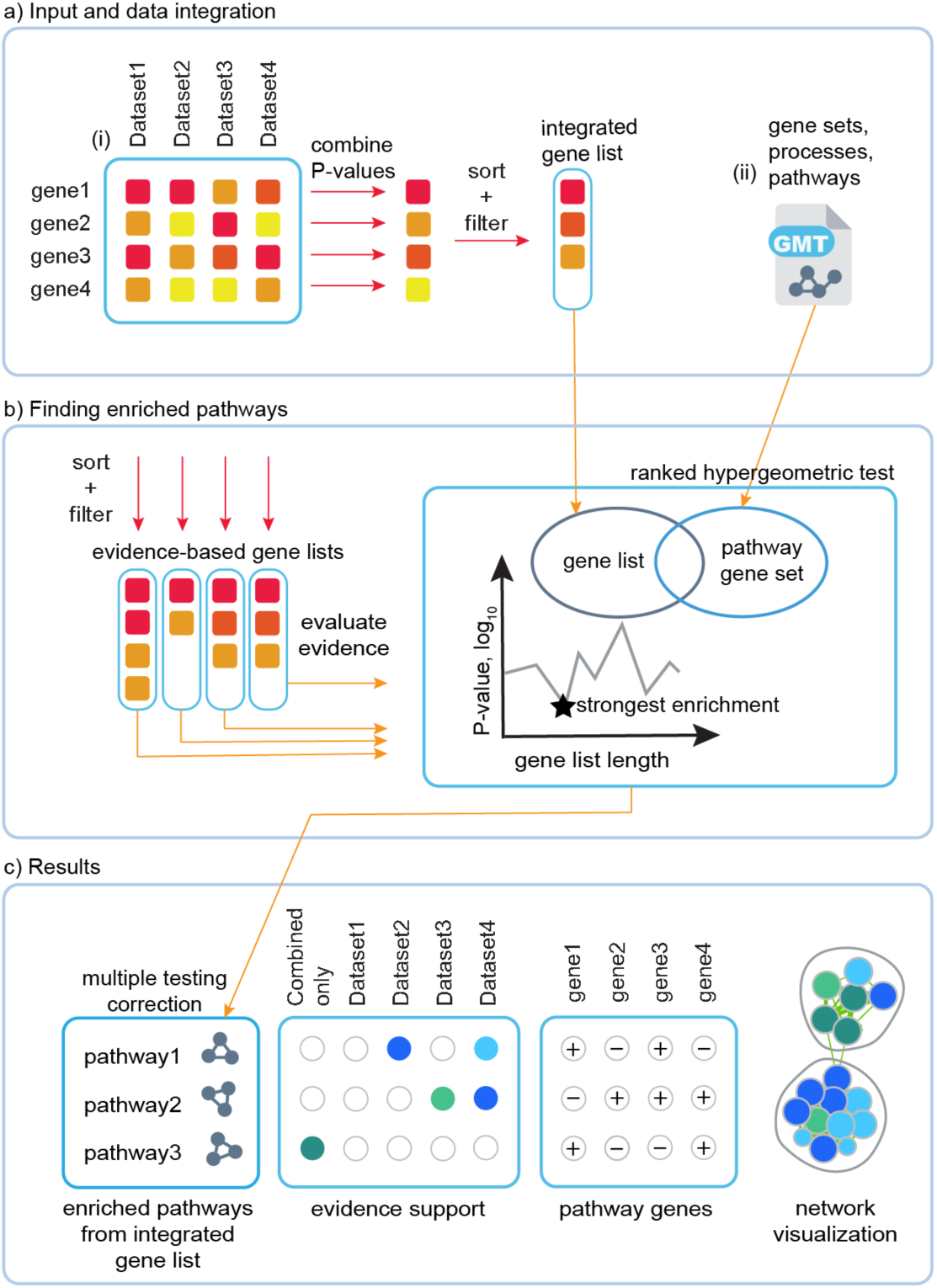
Method overview. **(a)** ActivePathways requires as input (i) a matrix of gene P-values for different omics datasets, and (ii) a collection of gene sets corresponding to biological pathways and processes. Gene p-values are merged and filtered to produce an integrated gene list that combines evidence from omics datasets and is ranked by decreasing significance with a lenient threshold. **(b)** Pathway enrichment analysis is conducted on the integrated gene list as well as lists from individual omics datasets using the ranked hypergeometric test that determines the optimal level of enrichment in the ranked gene sub-list for every pathway. **(c)** Pathways enriched in the integrated gene list are corrected for multiple testing and significant findings are reported as results. Pathways enriched in individual omics datasets are labelled by supporting evidence (colored nodes), and pathways only enriched in the integrated gene list are highlighted separately. Pathway genes with significant signals in different omics data are also shown. Finally, datasets of enriched pathways provided by ActivePathways are visualized as enrichment maps in Cytoscape where nodes correspond to pathways and pathways with many shared genes are connected into networks representing broader biological themes.

### Pathway analysis of coding and non-coding mutations in 2,500 whole cancer genomes

We performed integrative pathway analysis of coding and non-coding driver predictions across 29 cancer patient cohorts of histological tumor types and 18 meta-cohorts combining multiple types of tumors, with 47 cohorts in total (**Supplementary Table 1**). ActivePathways found at least one significantly enriched process or pathway in the majority of these cohorts (42/47 or 89%, *Q*_*pathway*_<0.05) (**Figure 2a**). We analyzed the omics evidence supporting predictions of enriched pathways and found that most cohorts showed enrichments in pathways supported by protein-coding driver scores of genes (37/47 or 79%). This serves as a positive control since the majority of currently known cancer driver genes have frequent protein-coding mutations.

**Figure 2.**
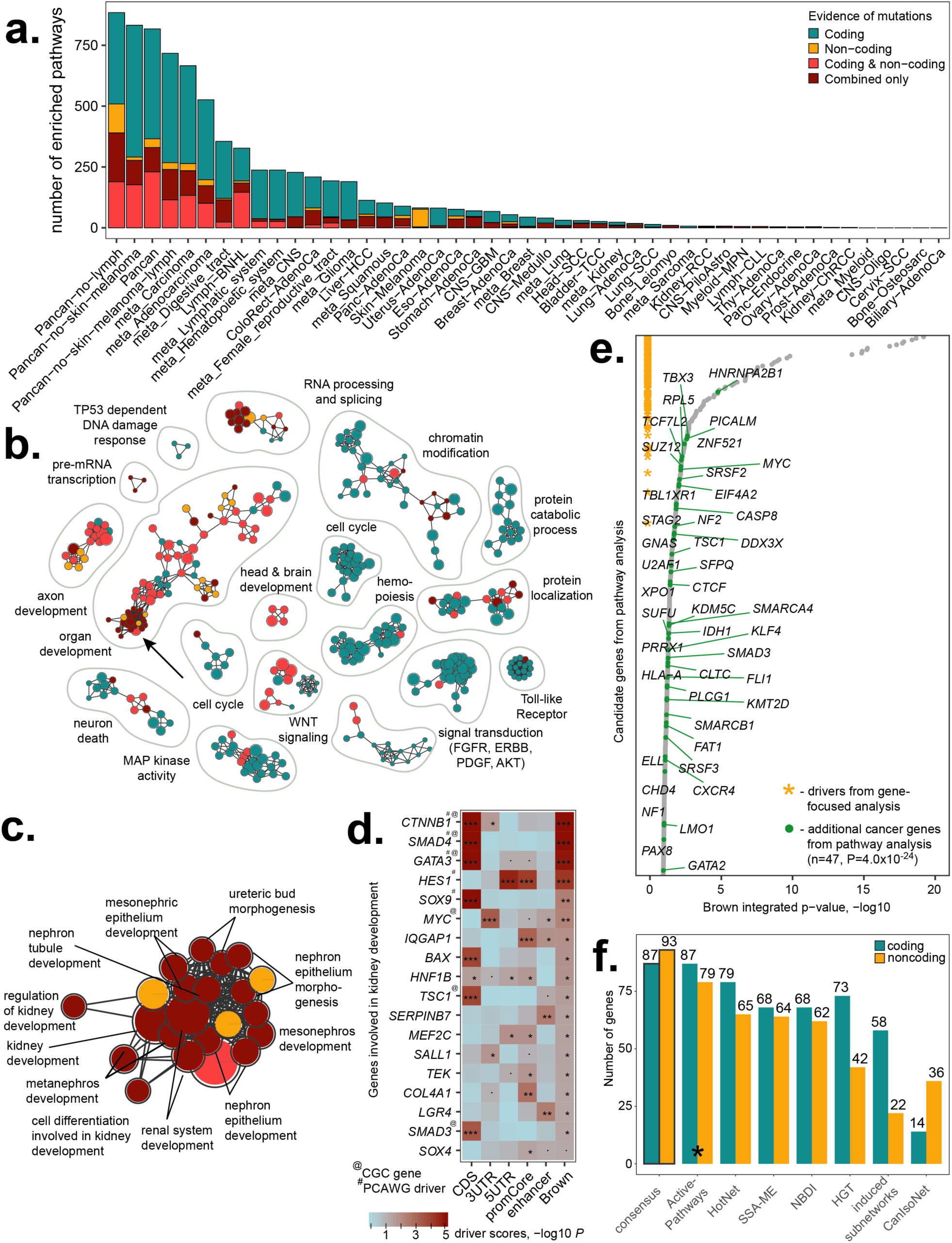
Pathway enrichment analysis of cancer driver genes with ActivePathways. **(a)** We analyzed consensus driver genes with frequent somatic mutations by integrating mutation scores of protein-coding and non-coding sequences (promoters, enhancers, and untranslated regions) across 47 cohorts of cancer patients with whole genome sequencing data from tumors. Bar plot shows number of significantly enriched pathways (Q<0.05) stratified by supporting evidence from driver predictions. The majority of pathways detected by ActivePathways are supported by protein-coding mutations, as expected (dark green bars), while non-coding mutations (orange, red) reveal additional pathways. Pathways shown in dark red are found only in the integrated gene list of coding and non-coding mutations but not in gene lists of individual mutation scores. **(b)** Enrichment map shows groups of statistically significant pathways characteristic of mutated genes in the adenocarcinoma cohort of 1,773 tumors. Nodes in the network diagram represent pathways that are connected with edges if the pathways are similar and share many genes. Groups of similar pathways were annotated manually. Nodes are colored by supporting evidence from coding and non-coding cancer mutations. **(c)** The group of enriched kidney developmental processes is apparent from integrated evidence of coding and non-coding mutations but is not found among coding or non-coding candidate genes separately (indicated with arrow in enrichment map). **(d)** P-value heatmap shows driver scores of genes involved in kidney developmental processes ranked by combined p-values of the integrated gene list (rightmost column). Top genes are expectedly detected as significantly mutated driver genes in the PCAWG consensus list while additional pathway-derived genes of the long tail of infrequent mutations are highlighted as well. Genes listed in the Cancer Gene Census (CGC) database are indicated with @ -symbol. **(e)** Integrated list of adenocarcinoma candidate driver genes used in the pathway enrichment analysis includes the majority of driver genes detected in the gene-focused consensus analysis by PCAWG (orange asterisks) and a long tail of infrequently mutated genes ranked by decreasing significance. Additional known cancer genes detected in the pathway analysis are indicated with green dots and occur more frequently than expected from chance alone. **(f)** Comparison of ActivePathways with six additional pathway and network analysis methods used in the PCAWG project. ActivePathways best recovers the consensus lists of pathway-implicated driver (PID) genes with coding and non-coding mutations. The consensus lists are shown in the leftmost bars of the plot and have been compiled through a majority vote of the seven methods in the PCAWG pathway and network analysis working group.

Non-coding mutations in genes also contributed to the discovery of frequently mutated biological processes and pathways: 24/47 cohorts (51%) showed significantly enriched pathways that were apparent when only analyzing non-coding driver scores separately for UTRs, promoters or enhancers. The majority of cohorts (41/47 or 87%) revealed enriched pathways that were apparent in the integrated gene list but not in any gene lists ranked by element-specific driver scores, emphasizing the value of our integrative approach. As expected, cohorts with more patient tumor samples generated more significantly enriched pathways (Spearman *ρ*=0.74, *P*=2.3x10^-9^; **Supplementary Figure 1**), suggesting that larger datasets are better powered to distinguish rarely mutated genes involved in biological pathways and processes. Discovery of pathways enriched in non-coding mutations suggests that pathway analysis is an attractive strategy for illuminating the dark matter of the non-coding cancer genome.

We studied the adenocarcinoma meta-cohort with 1,773 samples of 16 tumor types whose integrated list of 432 candidate genes (unadjusted *P*_*gene*_<0.1) associated with 526 significantly enriched pathways (*Q*_*pathway*_<0.05) (**Figure 2b**). As expected, the majority of pathways were only supported by genes with frequent coding mutations (328/526 or 62%). However, 101 pathways were supported by both coding and non-coding gene mutations, 72 were only apparent in the integrated analysis of all evidence, and 25 were only found among genes with significant non-coding mutations, thus expanding the set of candidate driver mutations in the non-coding cancer genome and demonstrating the value of integrative pathway analysis.

The major biological themes with frequent protein-coding mutations included hallmark cancer processes like *apoptotic signaling pathway* (24 genes; *Q*_*pathway*_=4.3x10^-5^) and *mitotic cell cycle* (8 genes; *Q*_*pathway*_=0.0026), and additional biological processes such as chromatin modification and RNA splicing that are increasingly recognized in cancer biology. Thus, our method captures the expected cancer pathways among driver genes with protein-coding mutations as positive controls. In contrast to these solely protein-coding driver associations, a large group of developmental processes and signal transduction pathways was enriched in genes with coding as well as non-coding mutations; for example *embryo development process* was supported by mutations in exons, 3’UTRs and gene promoters (68 genes; *Q*_*pathway*_=2.9x10^-12^), while *repression of WNT target genes* was only apparent in the integrated analysis of coding and non-coding mutations but not in either alone (5 genes, *Q*_*pathway*_=0.016; REAC:4641265). Thus, our method evaluates contribution of omics evidence towards pathway enrichments and finds additional associations that are not apparent in any provided dataset.

### ActivePathways highlights pathway-associated cancer genes in the long tail of infrequent non-coding mutations

We focused on a group of processes involved in kidney development that were only detected in the integrated analysis (**Figure 2c-d**). ActivePathways found 18 genes involved in these processes, only five of which were predicted as driver genes in the consensus driver analysis of the PCAWG project^14^. Additional known cancer genes included the oncogene *MYC* with 13 patients with 3’UTR mutations (*P*_*UTR3*_=4.8x10^-4^; *Q*_*UTR3*_=0.42), the transcription factor *SMAD3* of the TGF-β pathway with 14 patients with protein-coding mutations (*P*_*CDS*_=4.0x10^-4^; *Q*_*CDS*_=0.37) and the growth inhibitory tumor suppressor gene *TSC1* with 23 patients with protein-coding mutations (*P*_*CDS*_=1.4x10^-4^; *Q*_*CDS*_=0.17) as well as candidate cancer genes such as *IQGAP1* with 10 patients with promoter mutations (*P*_*promoter*_=8.2x10^-4^; *Q*_*promoter*_=0.62) that encodes a signaling protein that regulates cell motility and morphology. The additional genes remained below the FDR-adjusted significance cut-off in the gene-focused consensus driver analysis, however were found by ActivePathways due to pathway associations with frequently mutated developmental genes. These results highlight the potential of our method to find known and candidate cancer genes with rare coding and non-coding driver mutations through pathway-driven data integration.

We evaluated 333 candidate driver genes from the pathway analysis of the adenocarcinoma cohort (**Figure 2e**). These included as positive controls 60/64 significantly mutated genes identified in the PCAWG consensus driver analysis^14^, and an additional 47 genes of the COSMIC Cancer Gene Census database^12^, significantly more than expected by chance alone (seven genes expected, Fisher’s exact *P*=4.0x10^-24^), including *MYC, IDH1, NF1,* and *BCL9*. Additional genes were detected for several reasons. First, the integrated gene list was filtered using a lenient statistical cut-off (*P*_*gene*_<0.1) compared to a more stringent gene-focused driver analysis (*Q*_*gene*_<0.05). This resulted in 273/333 pathway-associated genes of the long tail that remainedbelow the significance threshold in the driver analysis. Second, the integration procedure combined multiple weaker p-values (coding regions, promoters, UTRs, enhancers) to a single stronger p-value for 17/333 pathway-associated genes including six cancer genes (*HNRNPA2B1*, *STAG2*, *TCF7L2*, *SUZ12*, *CLTC*, *ZNF521*) and improved the overall ranking of 220/333 genes among the input data, better explaining their membership in pathways and processes. However, a majority of all genes showed reduced significance after the integration procedure and were excluded from the pathway analysis, as the Brown combined p-value remained below the significance cut-off compared to any individual p-values of mutations in coding and non-coding regions of genes (3,112/3,543 or 88% genes with unadjusted *min(P*_*gene*_)<0.1 showed unadjusted Brown *P*_*gene*_>0.1). Fourth, the evidence evaluation step of the method identified pathway enrichments in gene lists ranked by individual sources of evidence and highlighted additional genes that did not pass significance cut-offs of the integration procedure. Thus, ActivePathways finds additional cancer genes in the long tail of mutations that are highlighted due to their pathway associations but remain below the significance cut-off in the gene-by-gene analysis.

### Benchmarking demonstrates the robustness and sensitivity of ActivePathways

We carefully benchmarked ActivePathways using multiple approaches. First, we compared its performance with six diverse methods used in the PCAWG pathway and network analysis working group^20^ (Hierarchical HotNet^21,22^, SSA-ME^23^, NBDI^24^, induced subnetwork analysis^22^, CanIsoNet^[*Kahraman*^ ^*et*^ ^*al,* *in*^ ^*prep*]^, and hypergeometric test). The methods used molecular pathway and network information to analyze the PCAWG dataset of predicted cancer driver genes^14^, and a subsequent consensus procedure derived pathway-implicated driver (PID) gene lists with coding (PID-C) and non-coding (PID-N) mutations based on a majority vote. Our method recovered PID-C and PID-N gene lists with the highest accuracy: 100% of coding driver genes (87/87) and 85% of non-coding candidates (79/93) were detected (**Figure 2f**).

We evaluated the robustness of ActivePathways to parameter variations and missing data. We varied the parameter *P*_*gene*_ that determines the ranked gene lists used in the pathway enrichment analysis (default threshold *P*gene<0.1). The majority of cohorts (40/47 or 85%) retrieved significantly enriched pathways even with a considerably more stringent threshold (*P*_*gene*_<0.001), however 67% fewer pathways were found compared to the default threshold in the median cohort (**Supplementary Figure 2**). We then evaluated the robustness of ActivePathways to missing data by randomly removing subsets of driver scores from the initial dataset. Even when removing 50% of gene driver scores with *P*<0.001, the majority of cohorts (37/47 or 79%) were found to have at least one significantly enriched pathway however 66% fewer pathways were found on average (**Supplementary Figure 3**).

**Figure 3.**
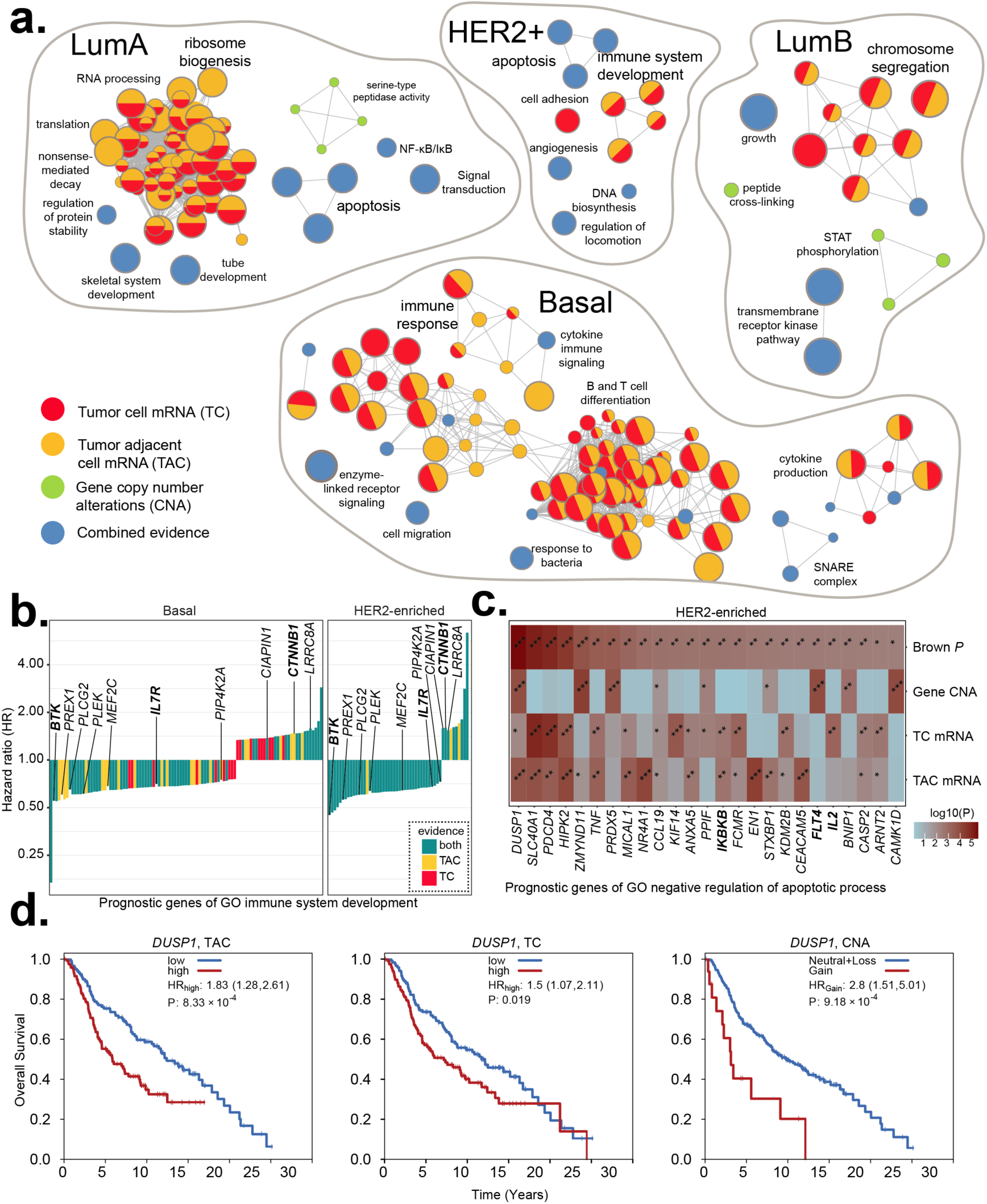
Prognosis-associated pathways in four molecular subtypes of breast cancer. (a) Enrichment maps of prognostic pathways and processes were found in an integrative analysis of mRNA abundance in tumor cells (TC), tumor-adjacent cells (TAC) and gene copy number alterations (CNA). Multi-colored nodes indicate pathways that were prognostic according to several types of molecular evidence. Blue nodes indicate pathways that were only apparent through merging of molecular signals. (b) Hazard ratios (HR) of prognostic genes of immune system development in basal and HER2-enriched subtypes of breast cancer. Strongest HR of TC, TAC is shown. Genes commonly found in basal and HER2-enriched tumors are shown. (c) Heatmap shows genes and corresponding p-values of the GO process “negative regulation of apoptotic process” found as prognostic in HER2-enriched breast cancer. Top row of the heatmap shows Brown p-values of merged evidence. (d) Kaplan-Meier plots show the strongest prognostic signal of the above apoptotic process associated with the DUSP1 encoding a protein phosphatase. DUSP1 significantly associates with reduced patient survival through increased tumor-adjacent mRNA level (left), increased tumor mRNA level (center) and gene copy number amplification (right). Known cancer genes are shown in boldface letters.

We tested ActivePathways with data simulations through 1,000 datasets for each of 47 patient cohorts and found no significant pathways in 92% of simulations (**Supplementary Figure 4**). Simulated data were obtained by randomly reassigning driver scores to different genomic elements, a conservative approach that disrupts gene and pathway annotations while retaining strong scores in the data. The median family-wise false discovery rate across cohorts (7.2%) slightly exceeded the applied multiple testing correction (*Q*<0.05). Higher rates were observed in cohorts including melanoma tumors, potentially due to abundant promoter mutations caused by impaired nucleotide excision repair in protein-bound genomic regions^25^. We evaluated quantile-quantile (QQ) plots of pathway-based p-values from ActivePathways and found that p-values from observed gene scores often deviated from the expected uniform distribution and appeared statistically inflated (**Supplementary Figure 5**). However, p-values derived from simulated gene scores showed no inflation in our simulations. Anticipating that the strongest cancer driver scores associate with protein-coding sequence, we studied datasets with simulated protein-coding gene scores and true non-coding scores. As expected, these partially simulated datasets expectedly showed less p-value inflation, suggesting that highly significant known cancer genes involved in many different pathways are responsible for inflation. Statistical testing of highly redundant pathways and processes violates the independence assumption of statistical tests and multiple testing procedures, a known caveat of pathway enrichment analysis^1,2^, which likely explains the observed distribution of significance values of our method.

**Figure 4.**
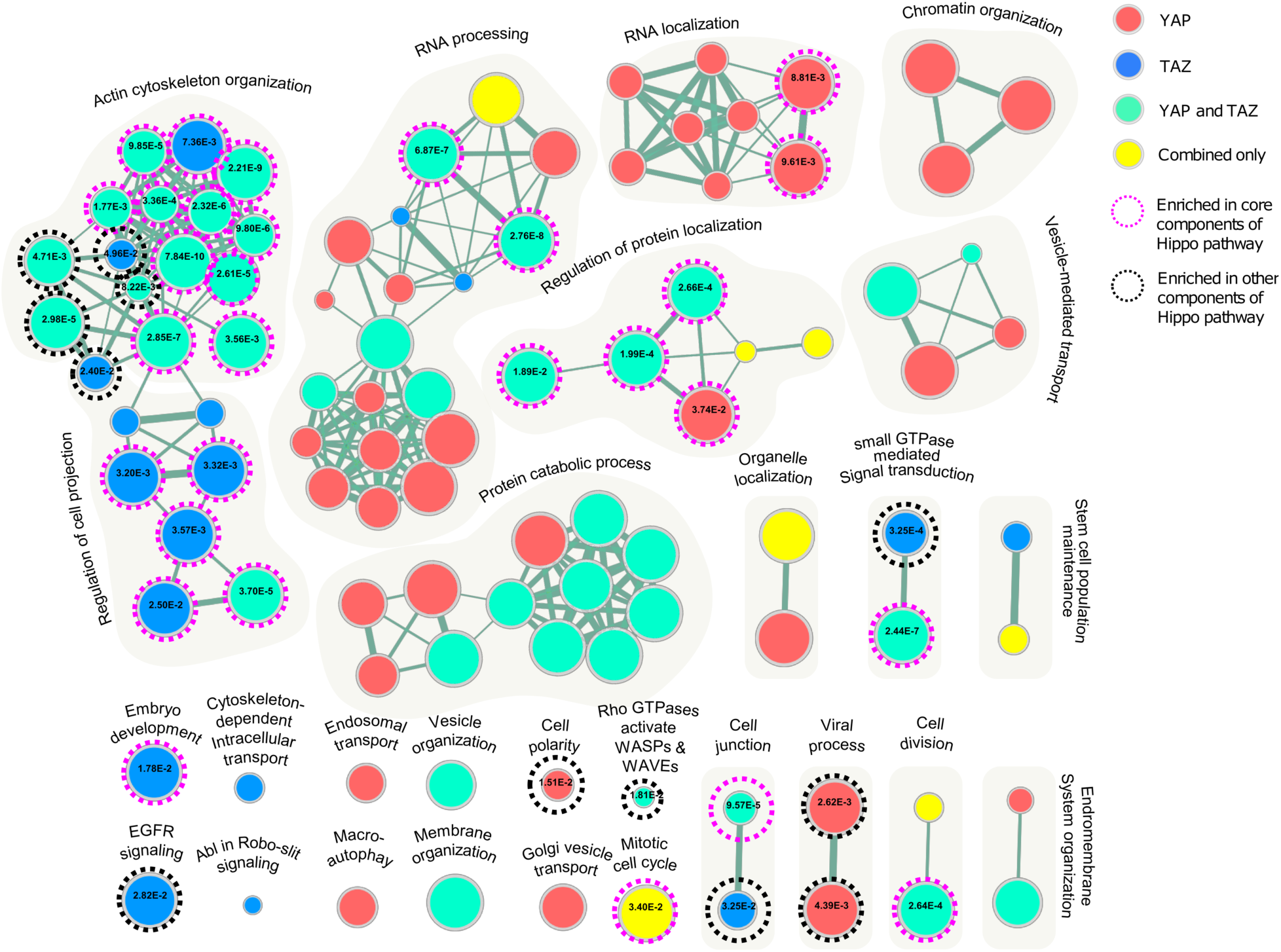
Pathway enrichment analysis of Hippo co-expression targets across human tissues. Enrichment map of pathways characteristic of genes co-expressed with transcription factors YAP and TAZ of the Hippo pathway across human tissues in the GTEx dataset. The Hippo pathway is involved in organ growth control and its predicted target genes are enriched in related biological processes and pathways. Nodes represent significantly enriched pathways that are colored by supporting evidence from co-expression targets of YAP or TAZ (red, blue), both transcription factors (green) or only the integrated list of target genes (yellow). We validated the detected pathways using a list of Hippo-related genes compiled from recent review papers and found that the majority of detected pathways included Hippo-related genes and 40% of pathways were enriched in these genes (indicated with dotted circles, enrichment p-values shown in nodes).

Collectively, these benchmarks show that ActivePathways is a sensitive and robust method for detecting significantly enriched pathways and processes through integrative analysis of multivariate omics data.

### Clinical analysis of genomic and transcriptional alterations of breast cancer subtypes reveals prognostic value of apoptotic, immune response and ribosomal genes

To demonstrate an integrative analysis of patient clinical information with multiple types of omics data, we then studied the pathways and processes associated with patient prognosis in breast cancer. We leveraged the METABRIC dataset^26^ using 1,780 breast cancer samples drawn from all four subtypes (HER2-enriched, basal-like, luminal-A, luminal-B) and evaluated all genes using three types of prognostic evidence. Gene expression profiles were deconvolved as mRNA abundance levels in tumor cells (TC) and tumor-adjacent cells (TAC) using the ISOpure algorithm^27^ and associated with these data with patient survival using median dichotomization and log-rank tests. Gene copy number alterations (CNA) were included as the third type of evidence and associated with patient survival using log-rank tests.

ActivePathways highlighted 192 significantly enriched GO biological processes and Reactome pathways across the four subtypes, of which nine were enriched in multiple subtypes and 33 were only apparent through the integrative pathway analysis but not in any omics evidence alone. Enrichment maps of significant results revealed immune response, apoptosis, ribosome biogenesis and chromosome segregation as the major groups of prognosis-associated pathways (**Figure 3a**).

Immune activity was associated with prognostic genes in basal-like and HER2-enriched breast cancers with significant enrichment of GO processes such as immune system development (*Q*_*basal*_=3.0x10^-4^, 113 genes; *Q*_*HER2*_=0.035, 61 genes) and lymphocyte differentiation (*Q*_*HER2*_=6.8x10^-4^, 46 genes; *Q*_*basal*_=8.4x10^-4^, 45 genes). The majority of genes of immune system development were associated with improved patient prognosis upon increased gene expression in tumor cells or tumor-adjacent cells, comprising 50/61 genes in the HER2-enriched subtype and 78/113 genes in the basal subtype (**Figure 3b**). Interestingly, only a minority of these genes (10) were significant in both of the two subtypes, suggesting different modes of immune activity in subtypes and emphasizing the power of our pathway-based approach. Basal-like breast cancers were associated with additional 67 terms involving immune response and blood cells, however no immune related terms were enriched for luminal subtypes of breast cancers. Prognostic features of immune-related genes in HER2-enriched and basal-like breast cancers are well known^28,29^. Our pathway-based findings indicate that immune activity in breast tumor cells and in the surrounding microenvironment negatively affect tumor progression and benefits the patient.

Apoptosis was associated with patient prognosis in HER2-enriched and luminal-A breast cancers through enriched GO processes such as negative regulation of apoptotic process (*Q*_*HER2*_=0.030, 122 genes; *Q*_*luminalA*_=0.015, 228 genes) and programmed cell death (*Q*_*HER2*_= 0.015, 125 genes; *Q*_*luminalA*_= 0.016, 231 genes) (**Figure 3c**). Anti-apoptotic pathways were only detected in the integrative analysis and not in genomic and transcriptomic gene signatures separately. Among the genes negatively regulating apoptosis, *DUSP1* provided the strongest prognostic signal in HER2-enriched breast cancers. This was apparent in the molecular stratification of samples by mRNA of tumor cells (log-rank *P*_*TC*_=0.019, HR=1.5) and tumor-adjacent cells (*P*_*TAC*_=8.3x10^-4^, HR=1.83) as well as gene copy number amplifications (*P*_*CNA*_=9.8x10^-4^, HR=2.8) (**Figure 3d**). *DUSP1* encodes a phosphatase signaling protein of the MAPK pathway that is over-expressed in malignant breast cancer cells and inhibits apoptotic signaling^31^. *HER2* over-expression is known to suppress apoptosis in breast cancer^30^. Anti-apoptotic signaling is a hallmark of cancer and expectedly associated with worse patient prognosis.

ActivePathways also identified prognostic pathway associations in single subtypes of breast cancer. For example, the prognostic genes for luminal-B subtype were enriched for chromosome segregation (*Q*_*luminalB*_=0.017, 41 genes) and related biological processes of GO. In agreement with this finding, problems with chromosome segregation have been associated with worse outcome in breast cancer^32^. As another example, luminal-A breast cancers were associated with prognosis in ribosomal and RNA processing genes, such as ribosome biogenesis (*Q*_*luminalA*_=6.9x10^-10^, 60 genes), and rRNA metabolic process (*Q*_*luminalA*_=1.8x10^-13^, 64 genes). Although not described specifically in the luminal-A subtype, ribosomal mRNA abundance has been shown to be prognostic in breast cancer as a marker of cell proliferation^33,34^. In summary, ActivePathways can be used for integrating clinical data with multi-omics information of molecular alterations. Such analyses can provide leads for functional studies and biomarker development.

### Co-expression analysis of Hippo master regulators across 54 human tissues recovers associated biological processes and genes

To study the use of ActivePathways in the context of healthy human tissues, we analyzed the dataset of 11,688 transcriptomes of 54 tissues from the GTEx project^5^, focusing on the Hippo signaling pathway involved in organ size control, tissue homeostasis and cancer^35,36^. We studied gene co-expression networks downstream of YAP and TAZ, the two master transcription factors of Hippo signaling, encoded by *YAP1* and *WWTR1*. YAP and TAZ are the evolutionarily conserved key effectors of the Hippo signaling in mammals. Inhibition of YAP/TAZ-mediated transcription regulates organ size control and tissue homeostasis in response to a wide range of intracellular and extracellular signals including cell-cell interactions, cell polarity, mechanical cues, ligands of G-protein-coupled receptors, and cellular energy status. We retrieved 2,117 putative Hippo transcriptional target genes that showed significant positive co-expression with either or both of the transcripts of *YAP* and *TAZ* across the human tissues in the GTEx dataset (*Q*_*gene*_<0.05). We used a robust rank aggregation method^37^ and retrieved transcriptional targets that were co-expressed with YAP or TAZ in a relatively large number of human tissues.

Analysis of the target genes using ActivePathways resulted in 101 significantly enriched pathways (*Q*_*pathway*_<0.05), including 39 supported by both sets of target genes, 37 supported by YAP1 targets, 18 supported by TAZ targets, and seven only apparent in the integrated list of target genes (**Figure 4**). The major biological themes of pathways and processes included regulation of cell polarity and cell junction, embryonic development, EGFR signaling, maintenance of stem cell population, actin cytoskeleton, and rho GTPase signaling that are all directly or indirectly related to Hippo signaling. We validated our analysis using 207 Hippo-related genes from review papers^35,36^ and confirmed that 83/101 pathways found by ActivePathways contained at least one of 59 Hippo-related genes, while 41 pathways were significantly enriched in Hippo-related genes (*Q*<0.05). However, the majority of genes documented in the literature (148/207) were not detected in the pathway analysis, potentially due to their post-transcriptional regulation or tissue-specific roles. Our analysis highlights known and candidate genes and pathways related to Hippo signaling and showcases the use of ActivePathways for functional analysis of transcription regulatory networks.

## Discussion

Integrative pathway enrichment analysis helps distill thousands of high-throughput measurements to a smaller number of pathways and biological themes that are most characteristic of the experimental data, ideally leading to mechanistic insights and novel candidate genes for follow-up studies. The primary advantage of our method is the fusion of gene significance across multiple omics datasets. This allows us to identify additional pathways and processes that are not apparent individually in any analyzed dataset. In our example of cancer driver discovery, pathway analysis is complementary to gene-focused driver discovery as it also focuses on sub-significant genes with coding and non-coding mutations clustered into known and novel biological processes of cancer. In the clinical analysis of breast cancer subtypes, we find prognostic genes and pathways active in tumor cells, the microenvironment, or both. A subset of these findings, such as anti-apoptotic signaling, is only apparent through data integration.

Our general pathway analysis strategy is applicable to diverse kinds of omics datasets where well-calibrated p-values are available for the entire set of genes or proteins. One may study a series of genomic, transcriptomic, or proteomic experiments or combine these into a multi-omics analysis. Data from epigenomic experiments and genome-wide association studies can be analyzed after genome-wide signals have been appropriately mapped to genes. Clinical and phenotypic information of patients can be also included through association and survival statistics. Our method is expected to work with unadjusted as well as multiple-testing adjusted p-values, however it is primarily intended for un-adjusted p-values for increased sensitivity. P-value adjustment for multiple testing is conducted at the pathway level rather than at a gene level. P-values from omics datasets are easier to interpret than raw signals as gene-based p-values are expected to account for experimental and computational biases specific to each analyzed dataset, while accounting for multi-omics factors comprehensively in a single generally applicable pathway-based model would be likely impossible. In our example of cancer driver discovery, appropriately computed p-values account for confounding factors of somatic mutations such as gene sequence length and nucleotide content, mutation signatures active in different types of tumors^38^ and biological cofactors of mutation frequency such as transcription and replication timing^39^, while pathway analysis of mutation counts or frequencies would maintain such biases in results.

Our analysis comes with important caveats. First, we only evaluate genes annotated in pathway databases that have variable coverage, rely on frequent data updates^40^ and may miss novel sparsely annotated candidate genes. The most general pathway enrichment analysis considers biological processes and molecular pathways however many kinds of gene sets available in resources such as MSigDB^41^ can be used to expand the scope of ActivePathways. Second, pathway information is highly redundant and analysis of rich *omics* datasets often results in many significant results reflecting the same underlying pathway. We address this redundancy by visualizing and summarizing pathway results as enrichment maps^2,19^ that help distill general biological themes comprised of multiple similar pathways and processes. Statistical inflation of results accompanied by biological redundancy is addressed by a stringent multiple testing correction. Third, the analysis treats pathways as gene sets and does not consider their interactions. This expands the scope of our analysis to a wider repertoire of pathways and processes as reliable mechanistic interactions are often context-specific and limited to a small subset of well-studied signaling pathways. Several methods such as HotNet^21^, PARADIGM^42^ and GeneMania^43^ model pathways and *omics* datasets through gene and protein interactions.

Translation of discoveries into improved human health through actionable mechanistic insights, biomarkers, and molecular therapies is a long-standing goal of biomedical research. Next-generation projects such as ICGC-ARGO (https://www.icgcargo.org/) aim to collect multi-*omics* datasets with detailed clinical profiles of patients and thus present novel challenges for pathway and network analysis techniques. In summary, ActivePathways is integrative pathway analysis method that improves systems-level understanding of cellular organization in health and disease.

## Methods

### Integrated and evidence-based gene lists

The main input of ActivePathways is a matrix of p-values where rows include all genes of a genome and columns correspond to omics datasets. To interpret multiple omics datasets, a combined p-value was computed for each gene using a data fusion approach, resulting in an integrated gene list. The integrated gene list was computed gene-by-gene by merging all p-values of a given gene into one combined p-value using the Brown’s extension^17^ of the Fisher’s combined probability test that accounts for overall co-variations of p-values from different sources of evidence. The integrated gene list of Brown p-values was ranked in order of decreasing significance and filtered using a lenient threshold of unadjusted *P*<0.1. Evidence-based gene lists representing different omics datasets were based on ranked P-values from individual columns of the input matrix, using the same significance threshold.

### Statistical enrichment of pathways

Statistical enrichment of pathways in significance-ranked lists of candidate genes was carried out with the ranked hypergeometric test. The test considered one pathway gene set at a time and analyzed increasing subsets of input genes from the top of the ranked gene list. The same procedure was used for integrated and evidence-based gene lists. At each iteration, the test computed the hypergeometric enrichment statistic and P-value for the set of genes shared by the pathway and top sub-list of the input gene list. For optimal processing speed, only gene lists ending with a pathway-related gene were considered as these most impact significance of enrichment. The ranked hypergeometric statistic selected the input gene sub-list that achieved the strongest enrichment and the smallest p-value as the final result for the given pathway, as

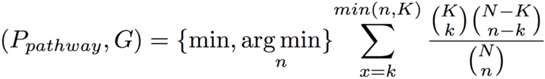

where *P*_*pathway*_ stands for the hypergeometric P-value of the pathway enrichment at the optimal sub-list of the significance-ranked candidate genes, *G* represents the length of the optimal sub-list, i.e. the number of top genes from the input gene list, *N* is the number of protein-coding genes with annotations in the pathway database, i.e., in Gene Ontology and Reactome, *K* is the total number of genes in a given pathway, *n* is the number of genes in a given gene sub-list considered, and *k* is the number of pathway genes in the considered sub-list. For a conservative estimate of pathway enrichment, we considered as background *N* the universe of genes contained in pathway databases and ontologies rather than the complete repertoire of protein-coding genes. To obtain candidate genes involved in the pathway of interest, we intersected pathway genes with the optimal sub-list of candidate genes. The ranked hypergeometric p-value was computed for all pathways and resulting p-values were corrected for multiple testing using the conservative Holm-Bonferroni family-wise error rate (FWER) method^18^. Significant pathways were reported (*Q*<0.05).

### Evaluating *omics* evidence of enriched pathways

The integrated gene list was analyzed the using ranked hypergeometric test and enriched pathways were reported as results. Each evidence-based gene list representing an omics dataset was also analyzed for enriched pathways with the ranked hypergeometric test. Pathways found in the integrated gene list were labelled for supporting evidence if they were also found as significant in any evidence-based gene list. A pathway was considered to be found only through data integration and labelled as *combined-only* if it was identified as enriched in the integrated gene list but was not identified as enriched in any of the evidence-based gene lists at equivalent significance cutoffs (*Q*<0.05). Each detected pathway was additionally annotated with pathway genes apparent in the optimal sub-list of candidate genes, separately for the integrated gene list and each evidence-based gene list.

#### Gene scores of cancer mutations

We analyzed p-values of genes reflecting their statistical significance as candidate cancer drivers for multiple cohorts of cancer patients with whole genome sequencing data. The scores were compiled in the driver discovery analysis of the PCAWG project as a consensus of multiple independent methods^14^. The input matrix of gene scores (*P*-values) included all protein-coding genes as rows and their genomic elements as columns (exons, 5’ and 3’ untranslated regions (UTRs), promoters, enhancers). Elements with missing p-values were assigned *P*=1. Genes with multiple enhancers were assigned the score of the most significant enhancer, and enhancers with more than five associated genes were excluded prior to selection.

#### Pathways and processes

We used gene sets corresponding to biological processes of Gene Ontology^15^ and molecular pathways of the Reactome database^16^ downloaded from the g:Profiler web server^9^. Large general gene sets with more than a thousand genes and small specific gene sets with less than five genes were excluded.

### Enrichment map visualization

ActivePathways provides input files for the EnrichmentMap app^19^ of Cytoscape^45^ for network visualization of similar pathways and their coloring according to supporting omics evidence. Enrichment maps for adenocarcinoma driver mutations, breast cancer prognostics, and Hippo transcriptional networks were visualized with stringent pathway similarity scores (Jaccard and overlap combined coefficient 0.6) and manually curated for the most representative groups of similar pathways and processes. Singleton pathways that were redundant with larger groups of pathways were discarded. Coloring of pathways in the adenocarcinoma enrichment map was rearranged by merging colors of pathways supported by non-coding mutation scores of promoters, enhancers and/or UTRs into one group.

### Analysis of coding and non-coding mutations of the PCAWG pan-cancer dataset

We used ActivePathways to analyze driver predictions of coding and non-coding mutations across >2,500 whole cancer genomes of the ICGC-TCGA PCAWG Project. P-values of driver predictions were computed separately for protein-coding sequences, promoters, enhancers and untranslated regions (UTR3, UTR5) in the PCAWG driver discovery study by Rheinbay *et al*^14^ across multiple subsets of samples representing histological tumor types and pan-cancer cohorts. We used gene-enhancer mapping predictions provided by PCAWG, excluded enhancers with more than five target genes, and selected the most significant enhancer for each gene, if any. Unadjusted p-values for coding sequences, promoters, enhancers and UTRs were compiled as input matrices and analyzed as described above. Missing p-values were interpreted as ones. Results from ActivePathways were validated with two lists of cancer genes. Predicted drivers from the gene-focused PCAWG driver analysis^14^ were selected as statistically significant findings (*Q*<0.05) following a stringent multiple testing correction spanning all types of elements (exons, UTRs, promoter, enhancers). The curated list of known cancer genes was retrieved from the COSMIC Cancer Gene Census (CGC) database^12^. One-tailed Fisher’s exact tests were used to estimate enrichment of these genes using all protein-coding genes as background.

### Analysis of prognostic genes in breast cancer

ActivePathways was used to evaluate prognostic pathways in breast cancer using multiple types of omics data. mRNA gene expression data and gene copy number alteration (CNA) data of the were derived from the METABRIC cohort of 1,991 patients with a single primary fresh frozen breast cancer specimen each^26^. Curtis *et al*^26^ classified the patients into the intrinsic breast cancer subtypes using the PAM50 mRNA-based classifier^44^ resulting in 330 basal-like breast cancers, 238 HER2-enriched breast cancers, 721 luminal-A breast cancers, 491 luminal-B breast cancers. Using these data, we computationally deconvolved tumor cell (TC) mRNA and tumor adjacent cell (TAC) mRNA abundance levels from the bulk profiled specimens. TC mRNA was deconvolved using ISOpure^27^ run on MATLAB release 2010b. TAC mRNA was computed using the ISOpure.calculate.tac function from the R package ISOpureR v1.1.2. ISOpure was run independently for each breast cancer subtype. The mRNA univariate survival analysis was conducted as follows. For each gene, patients were dichotomized based on mRNA abundance. Dichotomization was either based on the median mRNA abundance for that gene or a fixed value of 6.5. Based on the mRNA abundance distribution of genes on the Y chromosome in female samples, 6.5 was estimated as the threshold for noise for non-expressed genes. Median dichotomization was used if the median was above 6.5 or if there were no events in one of the groups when dichotomizing based on 6.5. The high and low mRNA abundance groups were compared by univariate log-rank tests for overall survival. TC and TAC mRNA abundance were evaluated independently. Survival modelling was performed in the R statistical environment (v3.4.3) using the survival package (v2.42-3). The CNA univariate survival analysis was conducted as follows. For each gene, we assessed whether more gains or losses were apparent. The copy number status with a higher count was subsequently used to separate patients into two groups: those with the chosen copy number status and the remaining patients. The two groups were then used for overall survival modelling with log-rank tests in the R statistical environment (v3.4.3) using the survival package (v2.42-3).

### Co-expression analysis of GTEx transcriptomes

The RNAseq dataset of human tissues was downloaded from GTEx v7 data portal (https://www.gtexportal.org/home/). The dataset included transcript abundance values of 21,518 protein-coding genes in 11,688 samples across 54 tissues. Tissues with less than 25 available samples and low gene expression (mean TPM<1.0) were excluded from further analysis. Positive pairwise Pearson correlations of gene expression values of *YAP* and *TAZ* (symbols *YAP1, WWTR1*) and their putative target genes were investigated in individual tissues and ranked by statistical significance of correlation tests. Tissue-specific ranked correlations of target genes were then integrated into two master lists of target genes of YAP and TAZ, respectively, reflecting target genes that were consistently positively co-regulated with corresponding transcripts across a significant subset of considered human tissues. We used the robust rank aggregation (RRA) method developed by Kolde *et al*^37^ and filtered co-expressed genes by significance using the default parameters of RRA (*Q*_*gene*_<0.05). Significantly enriched pathways among the putative target genes of YAP and TAZ were detected using ActivePathways. We validated the pathways by investigating their agreement with known Hippo-related genes from recent review papers^35,36^. We tested each pathway for enrichment of literature-derived Hippo genes using Fisher’s exact tests and filtered significant findings after multiple testing correction (Q<0.05).

### Method benchmarking

We benchmarked ActivePathways using multiple approaches, including simulated datasets, parameter variations, and partial replacement of strong scores with missing values. Benchmarking was carried out with the PCAWG dataset of coding and non-coding cancer driver predictions. To evaluate false discovery rates of ActivePathways, we created simulated datasets by randomly reassigning all observed driver scores to random genes and genomic elements. Simulations were conducted separately for different tumor cohorts. One thousand simulated datasets were analyzed with ActivePathways and those with at least one significantly detected pathway counted towards false discovery rates. Additional simulations maintained the positions of non-coding driver scores among gene scores and randomly reassigned protein-coding driver scores, expectedly leading to a reduction in detected pathways as the input datasets primarily included strong scores in protein-coding gene regions. Quantile-quantile analysis and QQ-plots were used to compare p-value distributions of pathways discovered from true driver scores, driver scores with shuffled driver scores, and driver scores shuffled entirely. To evaluate robustness of ActivePathways, we randomly replaced a fraction of significant driver p-values in input matrices (*P*<0.001) with insignificant p-values (*P*=1). We tested different fractions of missing values (10%, 25%, 50%) across a thousand datasets of driver scores with randomly selected missing data points and concluded that most cohorts included significantly enriched pathways even with large fractions of missing data. To further evaluate robustness, we tested different values of the Brown *P*-value threshold used to select the integrated gene list for pathway enrichment analysis. The default parameter value (*P*_*gene*_<0.1) was compared to alternative values (0.001, 0.01, 0.05, 0.2). We concluded that ActivePathways found enriched pathways in most tumor cohorts even at more stringent gene selection levels.

### Availability

ActivePathways is freely available as an R package and source code on the GitHub repository https://github.com/reimandlab/ActivePathways and the Comprehensive R Archive Network (CRAN).

## Acknowledgements

This work was funded by Ontario Institute for Cancer Research (OICR) Investigator Awards to J.R. and P.C.B. provided by the Government of Ontario; Operating Grant to J.R. from Cancer Research Society (CRS) (#21089); Natural Sciences and Engineering Research Council of Canada (NSERC) Discovery Grant to J.R. (#RGPIN-2016-06485), and the Canada First Research Excellence Fund, University of Toronto Medicine by Design to J.R. H.Z. was supported by a CIHR Canadian Graduate Scholarship. J.B. was supported by a BioTalent Canada Student Internship. P.C.B. was supported by TFRI and CIHR New Investigator Awards.

